# Development of a Bacterial Colorimetric Reporter System for Functional Screening of SARS-Cov-2 Main Protease Inhibitors Using Plant Preparations (juices): A Proof-of-Concept Study

**DOI:** 10.64898/2025.12.12.694002

**Authors:** Shaza S Issa, Andrew A Zelinsky, Haidar J Fayoud, Roman R Zhidkin, Tatiana V Matveeva

**Affiliations:** Department of Genetics and Biotechnology, St. Petersburg State University, 199034 St. Petersburg, Russia; Laboratory of Amyloid Biology, St. Petersburg State University, St. Petersburg 199034, Russia; Laboratory of Proteomics of supra-organizational Systems №7, All-Russia Research Institute for Agricultural Microbiology, 196608 St. Petersburg, Russia; Department of Ecological genetics, Center for Biological regulation of Pesticide Use, All-Russian Institute of Plant Protection, 196608, St. Petersburg, Russia

**Keywords:** SARS-CoV-2, Mpro inhibitors, screening system, *E. coli*, colorimetric assay, β-galactosidase, pomegranate juice

## Abstract

SARS-CoV-2 main protease (Mpro) is essential for viral polyprotein processing and represents a prime target for antiviral drug discovery. However, most available screening strategies rely on biochemical and computational approaches that lack the biological context of living cells, or costly mammalian-cell based models. Therefore, there remains a shortage of simple and biosafe cellular models enabling rapid, functional screening of potential Mpro inhibitors, particularly those derived from natural sources and in urgent situations such as the COVID-19 pandemic. In this study, a bacterial colorimetric reporter system was developed that directly links SARS-CoV-2 Mpro activity to β-galactosidase function in *Escherichia coli*. To the best of our knowledge, the developed system represents the first bacterial colorimetric model for direct monitoring of SARS-CoV-2 Mpro inhibition in living cells. The system enables real-time visual detection of protease inhibition on X-gal-containing medium and provides a cost-effective, biologically relevant, biosafe alternative to existing screening assays. Functional validation was performed using pomegranate juice as a representative natural inhibitor source. The system provides a simple, scalable, and biosafe platform for the primary screening of antiviral candidates, including phytochemicals, under standard laboratory conditions.

## 1. Introduction

The main protease of novel coronavirus (SARS-CoV-2 Mpro), also known as the 3C-like protease (3CLpro) or non-structural protein 5 (Nsp5), plays a key role in viral replication by cleaving polyproteins pp1ab and pp1a into functional proteins [1, 2]. Accordingly, it is considered a primary drug target in the search for antiviral agents against COVID-19. In addition to its importance in the viral life cycle, SARS-CoV-2 Mpro has no homologs in the human host and exhibits a distinct substrate specificity, recognizing and cleaving at the motif Leu-Gln↓(Ser, Ala, Gly), which minimizes the risk of off-target effects, toxicity, or cross-reactivity in human cells [3, 4]. SARS-CoV-2 Mpro is also highly conserved among coronaviruses, reducing the possibility that resistance mutations will affect the efficacy of its inhibitors, making these inhibitors promising candidates for broad-spectrum antiviral development [4].

During the COVID-19 pandemic, the urgent need to identify anti-SARS-CoV-2 drugs, coupled with the requirement for biosafety level 3 facilities, led most research on Mpro inhibitors to rely on *in silico* or *in vitro* biochemical approaches rather than cell-based or *in vivo* models [5–7]. Although these strategies are useful for identifying promising candidates, they have their limitations [8]. For instance, computational docking and molecular dynamics simulations can predict the binding affinity of a potential inhibitor, but do not confirm functional inhibition [9], and their accuracy depends heavily on the data source, necessitating further experimental validation [10]. Similarly, *in vitro* biochemical assays, using purified SARS-CoV-2 Mpro under artificial conditions, often fail to account for important cellular factors such as protein folding and stability, as they lack the biological intracellular context [11]. Therefore, while these approaches are valuable for early screening, they provide limited insight into inhibitor-protease interactions in living cells, whereas mammalian cell-based assays remain costly and require high biosafety levels.

Plants have long served as sources of therapeutic agents, and to this day tens of thousands of medicinal species remain underexplored, offering a vast reservoir for drug discovery [12]. Their secondary metabolites, such as flavonoids, alkaloids, terpenoids, and phenolics, exhibit significant structural diversity shaped by evolutionary adaptation [13]. This diversity grants broad biological activity, target selectivity, multitarget potential, and generally low toxicity [13, 14]. Several phytochemicals, such as quercetin, baicalein, and epigallocatechin gallate (EGCG), have demonstrated inhibitory effects against SARS-CoV-2 Mpro in biochemical and computational studies [5]. In recent years, specifically since COVID-19 pandemic, plants have also been increasingly recognized not only as sources of pharmacologically active compounds but as functional foods capable of contributing to disease prevention and immune support through their complex mixtures of bioactive metabolites. A growing body of research highlights the role of polyphenol-rich fruits, vegetables, and plant-based products in modulating antiviral, anti-inflammatory, and immunomodulatory responses, positioning functional foods as promising complementary resources for managing viral infections and highlighting the effect of diet and proper nutrition on response to infections [15–19]. Given their accessibility, sustainability, and established safety, plant-derived preparations/compounds represent valuable sources for the search for broad-spectrum antiviral agents. However, despite numerous computational and biochemical findings, their functional activity against SARS-CoV-2 Mpro remains largely unexplored under cellular conditions. The lack of simple, affordable, and biosafe cellular systems has impeded this stage of validation, as mammalian models are technically demanding and resource-intensive [5, 20, 21].

To address this gap, we developed a bacterial reporter-based colorimetric system that enables functional assessment of SARS-CoV-2 Mpro inhibition directly in living *Escherichia coli* cells. To the best of our knowledge, this represents the first bacterial colorimetric model designed for this purpose. The system monitors SARS-CoV-2 Mpro activity, or its inhibition by tested juices, through disruption of β-galactosidase activity, using a colorimetric readout on X-gal-containing medium. This study describes the system’s construction and following functional validation using pomegranate juice, a natural source of several bioactive compounds with reported antiviral properties. Compared to existing methods, the system offers several advantages including real-time monitoring of SARS-CoV-2 Mpro activity in a living context, a simple visual readout enabling rapid evaluation of tested crude preparations/compounds, and a cost-effective, scalable platform that could be applied in a standard laboratory setting.

## 2. Materials and Methods

### 2.1. De novo assembly and sequence verification of codon-optimized SARS-CoV-2 Mpro gene

The nucleotide sequence of the *SARS-CoV-2 Mpro* gene was obtained from the isolate SARS-CoV-2/human/USA/WI-CDC-ASC210624442/2021 (GenBank accession no. OM036858.1). The sequence was manually codon-optimized, based on *E. coli* codon usage bias. *De novo* assembly of the gene was performed using a PCR-based overlapping fragment synthesis strategy. The entire gene was divided into 35 partially overlapping fragments, which were gradually assembled through 16 consecutive PCR reactions. Oligonucleotides and primers for intermediate amplicons were designed using UGENE and Vector NTI software (Invitrogen). Amplifications were carried out using a commercial PCR mix containing Taq DNA polymerase (Thermo Fisher Scientific). The final product was analyzed by electrophoresis on a 1% agarose gel, and verified by Sanger sequencing with terminal primers. Any detected point mutations were corrected by site-directed mutagenesis.

### 2.2. Plasmid construction

The *E. coli* colorimetric reporter system was constructed using the high-copy plasmid pUC19 as the vector backbone. This plasmid was selected for its strong replication in *E. coli* and its functional *lacZα* gene, which enables β-galactosidase-mediated hydrolysis of X-gal into a blue chromogenic product. To enable Mpro-dependent color switching, a specific SARS-CoV-2 Mpro cleavage site (AAAACCAGCGCGGTGCTGCAGAGCGGCTTCCGCAAAATGGAG) was inserted within the *lacZα* coding sequence without disrupting its open reading frame. The insert length was maintained as a multiple of three nucleotides, and the flanking codons were preserved to prevent any amino acid substitutions or premature stop codons. This design ensured that β-galactosidase activity remained intact unless the cleavage site was cleaved by Mpro.

The assembled codon-optimized *Mpro* gene was placed under the control of the lac promoter/operator to allow co-regulated expression of both *Mpro* and *lacZα*, and terminated by a T7 terminator sequence. The resulting lacP/O-*Mpro*-T7 terminator cassette was inserted into the same pUC19 plasmid. All cloning steps were verified by restriction digestion and Sanger sequencing. Construction strategy is shown in Figure 1.

**Figure 1.**
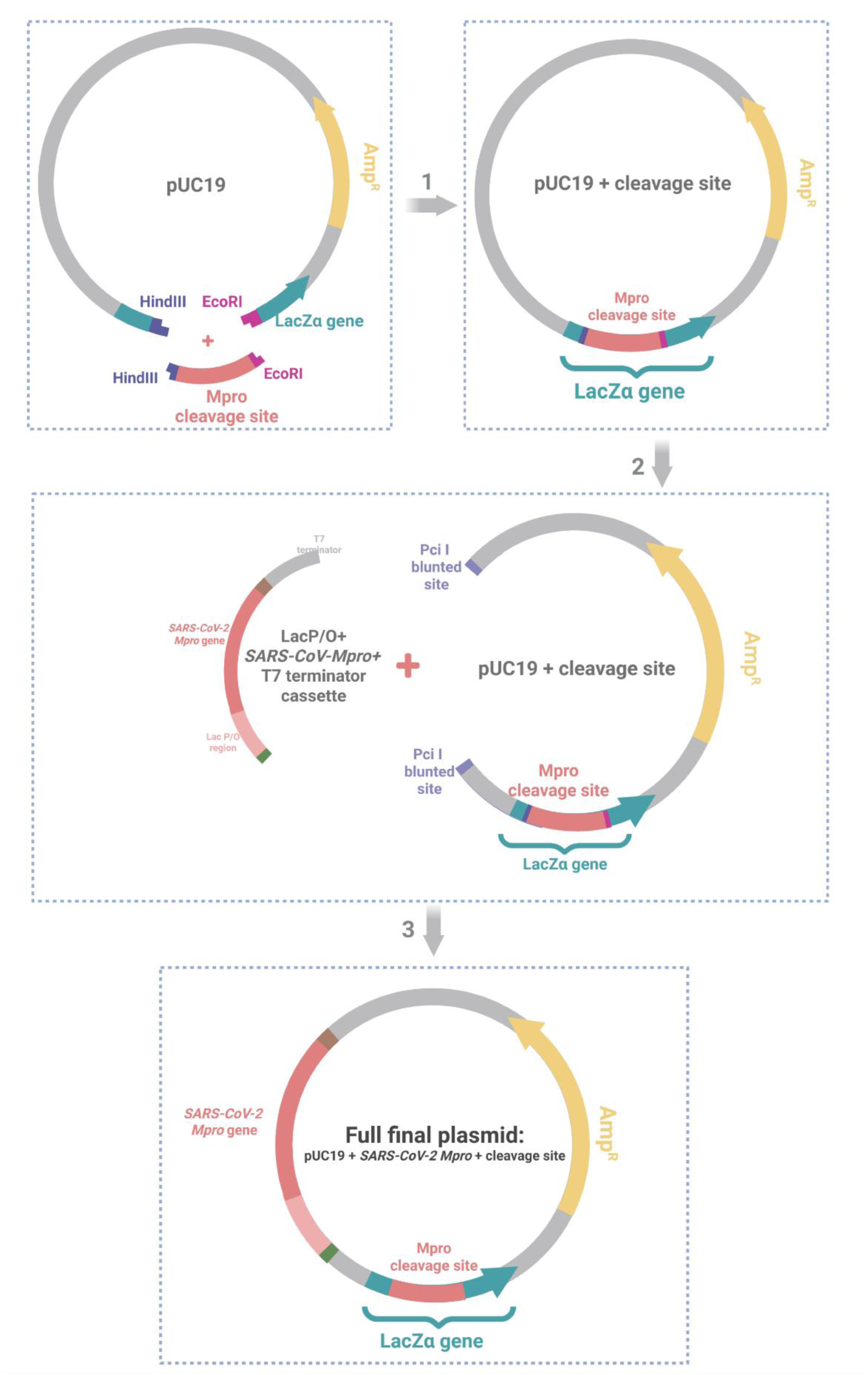
Schematic representation of the strategy used to obtain final construct.

### 2.3. Bacterial strain, transformation, and expression conditions

Chemically competent *E. coli* DH5α cells were used as the host strain for all cloning and expression experiments. This strain was selected for its high transformation efficiency, compatibility with high-copy-number plasmids such as pUC19, and reduced background protease and endonuclease activities, which support stable maintenance of recombinant constructs [22]. In addition, DH5α carries a chromosomal deletion that allows functional complementation with the *lacZα* fragment encoded by pUC19, making it ideal for colorimetric reporter assays based on β-galactosidase activity [22]. Transformation was performed by the standard heat-shock method, followed by recovery in LB medium and plating on LB agar supplemented with ampicillin (100 µg/mL). Plates were incubated at 37 °C for 16 h to obtain recombinant colonies. For expression experiments, single colonies were inoculated into 5 mL of LB broth containing ampicillin and grown overnight (∼16 h) at 37 °C with shaking at 250 rpm. The following day, cultures were refreshed into fresh LB medium (1:100 dilution) to maintain exponential growth. Cell density was monitored spectrophotometrically at 600 nm using a Hitachi U-2900 UV–Visible spectrophotometer. When the optical density (OD_600_) reached 0.6–0.9, expression was induced by adding isopropyl β-D-1-thiogalactopyranoside (IPTG; Helicon, Russia) to a final concentration of 0.2 mM. At the same time, 5-bromo-4-chloro-3-indolyl β-D-galactopyranoside (X-gal; dissolved in DMSO, SibEnzyme, Russia) was added to a final concentration of 0.2 mg/mL to enable colorimetric visualization of lacZ activity. Cultures were subsequently incubated at 26 °C with shaking (250 rpm) for an additional 16–18 h to allow color development.

### 2.4. SDS–PAGE analysis of Mpro expression

Cell lysates from *E. coli* DH5α carrying the recombinant pUC19–*Mpro* construct, and from control cells containing the empty pUC19 vector, were analyzed by SDS–PAGE. Prior to lysis, cell cultures were normalized by OD_600_ to ensure comparable protein loading. Samples were mixed with Laemmli SDS sample buffer containing 2% SDS (Bio-Rad, USA), boiled for 5 min, and loaded onto a 12% polyacrylamide gel together with a Kaleidoscope™ Prestained Protein Standard (Bio-Rad, USA) as a molecular weight marker. Electrophoresis was carried out using standard protocols. Proteins were stained with Coomassie Brilliant Blue (Bio-Rad, USA) and visualized with a Bio-Rad ChemiDoc™ imaging system and Image Lab 6.1 software.

### 2.5. Preparation of plant juices

Fresh pomegranates (*Punica granatum*) (local variety not specified) and fresh guelder rose berries (*Viburnum opulus*; калина) were obtained from a local market during their natural season (mid-November). All manipulations involving the juices were performed under sterile conditions. Approximately 25 g of pomegranate arils and 25 g of guelder rose berries were manually homogenized separately using a sterile mortar and pestle to obtain raw juices. The homogenates were clarified by centrifugation at 4200 rpm for 15 min at 4 °C, and the clear supernatants were collected. The pH of the supernatants was measured and adjusted to 4.0–4.5 with 10 N NaOH to reduce excessive acidity while preserving the potential contribution of their organic acids to the anti-Mpro activity. The juices were shortly stored at 4 °C until use.

### 2.6. Design and functional basis of the colorimetric screening system

The colorimetric screening system was designed to function as a gain-of-signal assay. It is based on the functional relationship between Mpro and the *lacZ* reporter gene. In the absence of Mpro, cells expressing *lacZ* alone produce a blue coloration in LB medium supplemented with X-gal and IPTG, due to β-galactosidase-mediated hydrolysis of X-gal yielding 5-bromo-4-chloro-indole, which oxidizes and dimerizes into a blue product. However, co-expression of Mpro leads to cleavage of its recognition site inserted within the *lacZα* fragment, disrupting β-galactosidase activity and preventing color formation. The addition of an inhibitory compound blocks Mpro activity, restoring β-galactosidase function and producing a gain of blue signal. The assay principle and expected color outcomes are illustrated in Figure 2.

**Figure 2.**
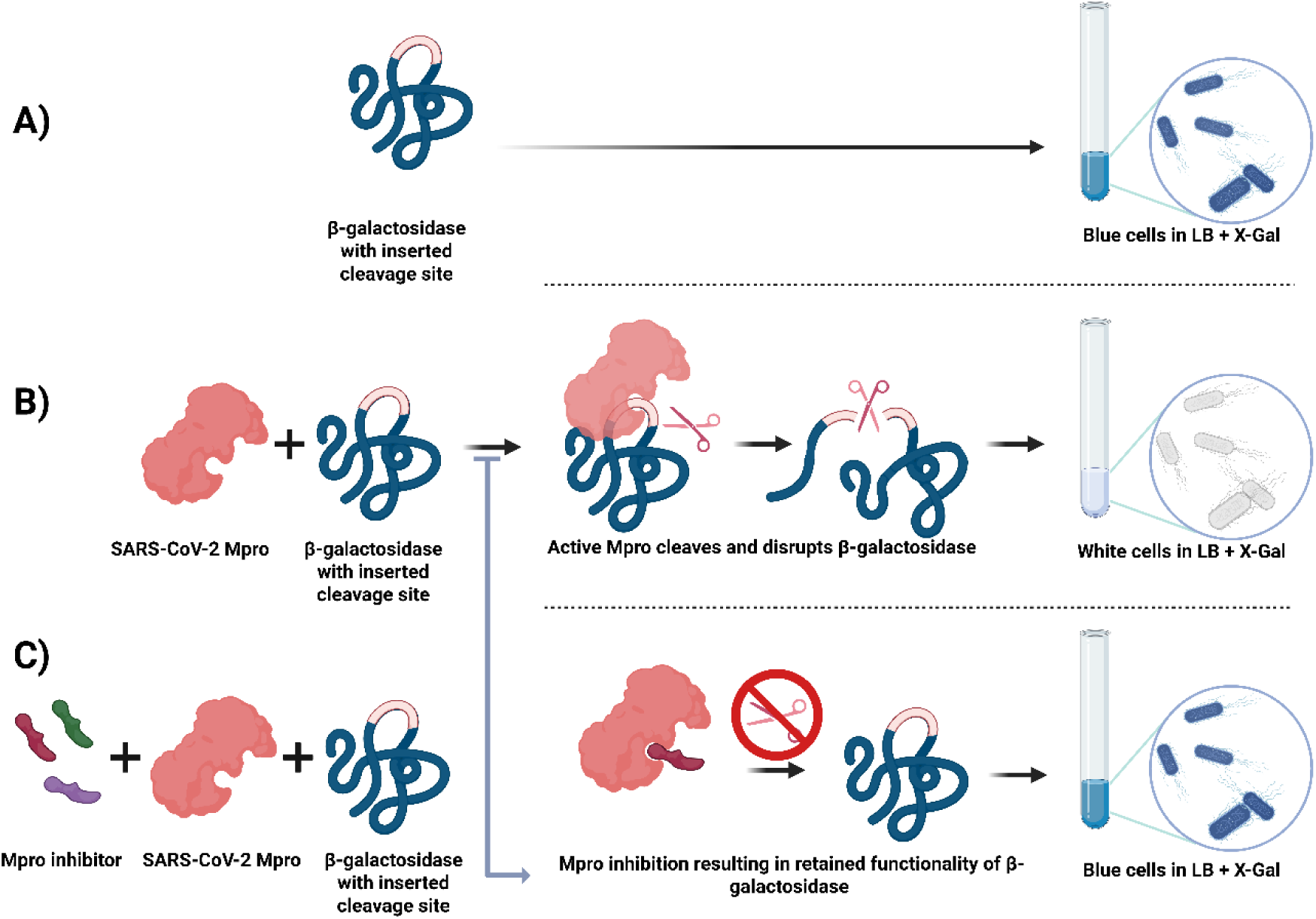
A schematic representation explaining the functional basis of the colorimetric screening system and the expected colorimetric outcomes in an X-gal-containing medium: A) when cells are expressing only the modified reporter, B) when cells are co-expressing the modified reporter and Mpro, and C) when cells are co-expressing the modified reporter and Mpro in the presence of an Mpro inhibitor.

### 2.7. Assay setup and inhibitor treatment (plant juices)

Freshly prepared, slightly pH-adjusted juices were tested at final concentrations of 2%, 5%, 10%, and 20% (v/v). All bacterial cultures were standardized to an identical OD_600_ of 0.8 prior to treatment (as achieved by refreshing overnight cultures, see subsection 2.3.). Juices were added at the time of induction simultaneously with IPTG and X-gal, and cultures were incubated at 26 °C for 16-18 h with shaking at 250 rpm to allow color development. To enable the use of high juice fractions (≥ 10% v/v) without diluting the cell density, OD-standardized cultures were centrifuged (2500 rpm, 2 min), supernatants discarded, and pellets immediately resuspended to 1.00 mL with a premix adjusted to yield a final 1X LB composition at the target juice fraction. Antibiotic, IPTG, and X-gal were added at the time of resuspension. All experimental groups were handled identically to maintain equal initial OD and consistent medium composition across treatments.

A defined set of internal controls was included in the assay to ensure the validity and interpretability of the colorimetric screening system, accounting for assay responsiveness, baseline β-galactosidase activity, and potential sources of experimental variability. The detailed composition and purpose of control groups are summarized in Table 1.

**Table 1.**
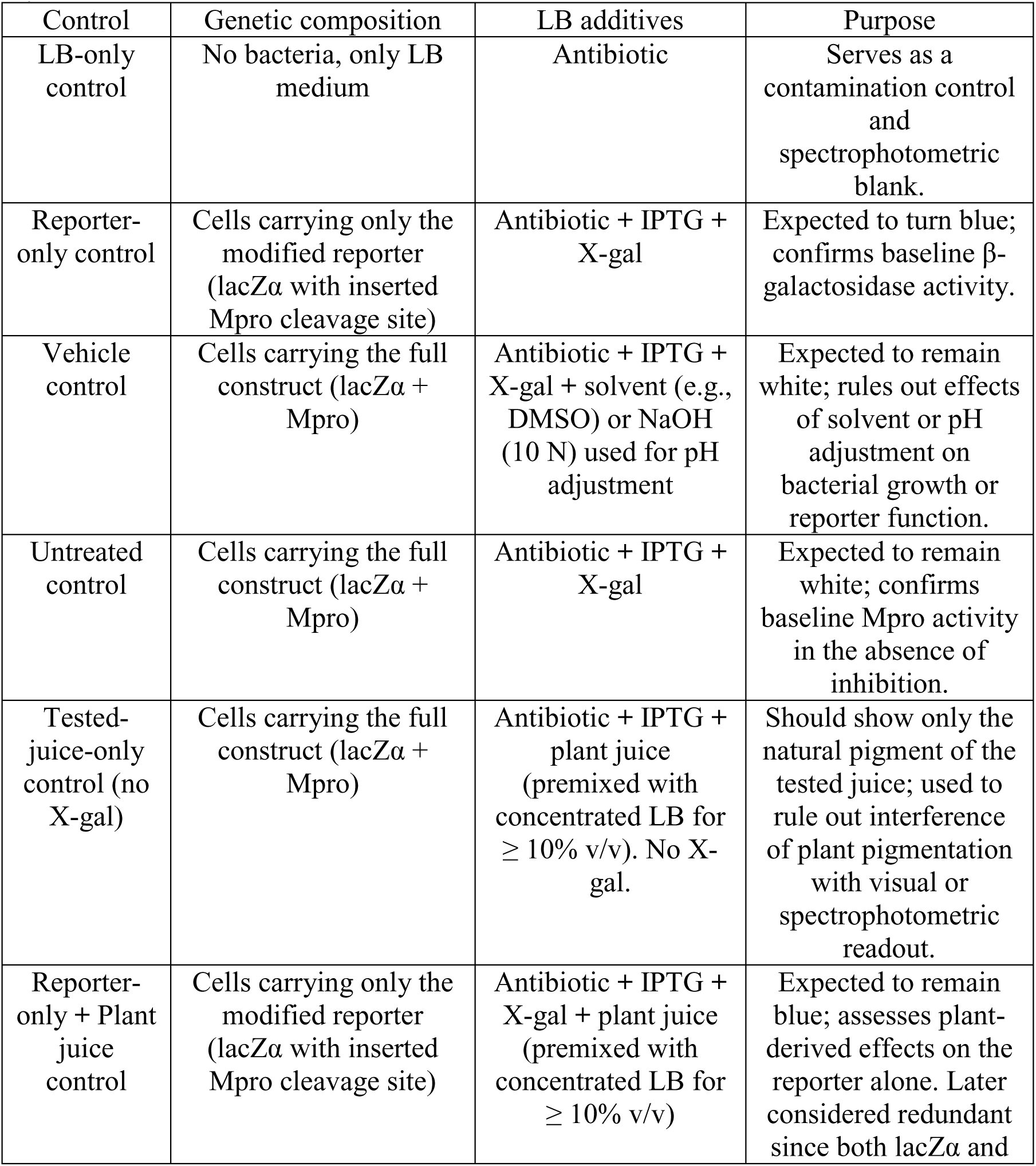
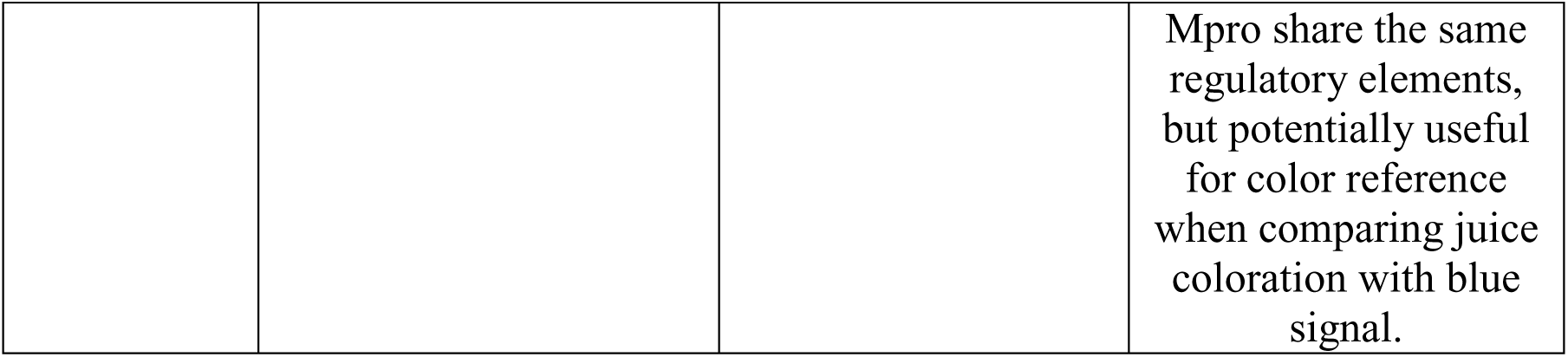
A summary of internal controls used in the developed colorimetric reporter system.

| Control | Genetic composition | LB additives | Purpose |
| --- | --- | --- | --- |
| LB-only control | No bacteria, only LB medium | Antibiotic | Serves as a contamination control and spectrophotometric blank. |
| Reporter-only control | Cells carrying only the modified reporter (lacZα with inserted Mpro cleavage site) | Antibiotic + IPTG + X-gal | Expected to turn blue; confirms baseline β-galactosidase activity. |
| Vehicle control | Cells carrying the full construct (lacZα + Mpro) | Antibiotic + IPTG + X-gal + solvent (e.g., DMSO) or NaOH (10 N) used for pH adjustment | Expected to remain white; rules out effects of solvent or pH adjustment on bacterial growth or reporter function. |
| Untreated control | Cells carrying the full construct (lacZα + Mpro) | Antibiotic + IPTG + X-gal | Expected to remain white; confirms baseline Mpro activity in the absence of inhibition. |
| Tested-juice-only control (no X-gal) | Cells carrying the full construct (lacZα + Mpro) | Antibiotic + IPTG + plant juice (premixed with concentrated LB for $\geq 10\%$ v/v). No X-gal. | Should show only the natural pigment of the tested juice; used to rule out interference of plant pigmentation with visual or spectrophotometric readout. |
| Reporter-only + Plant juice control | Cells carrying only the modified reporter (lacZα with inserted Mpro cleavage site) | Antibiotic + IPTG + X-gal + plant juice (premixed with concentrated LB for $\geq 10\%$ v/v) | Expected to remain blue; assesses plant-derived effects on the reporter alone. Later considered redundant since both lacZα and |
|  |  |  | Mpro share the same regulatory elements, but potentially useful for color reference when comparing juice coloration with blue signal. |

Cell growth inhibition by the tested juices was evaluated by comparing OD_600_ values measured immediately before and after incubation with the respective concentrations.

### 2.8. Quantification of color development

Following overnight incubation with the tested juices, bacterial cells were harvested by centrifugation at 4200 rpm for 15 min at 4 °C. The resulting pellets were resuspended in lysis buffer (8 M urea, 1.5 M Tris-HCl, pH 8.0), added at a ratio of 15 volumes of lysis buffer per gram of wet pellet, and incubated with gentle rotation at room temperature for 30 min. To ensure complete release of intracellular pigment, cell suspensions were subjected to ultrasonic disruption using a Q700 sonicator (Qsonica, USA) under the following conditions: 50% amplitude, 1 min on / 2 min off cycles. The lysates were then centrifuged at 3600 rpm for 5 min to remove cell debris, and the clarified supernatants were analyzed spectrophotometrically. The intensity of the developed blue chromogen was quantified by measuring absorbance at 650 nm. A schematic representation of the experimental workflow is shown in Figure 3.

**Figure 3.**
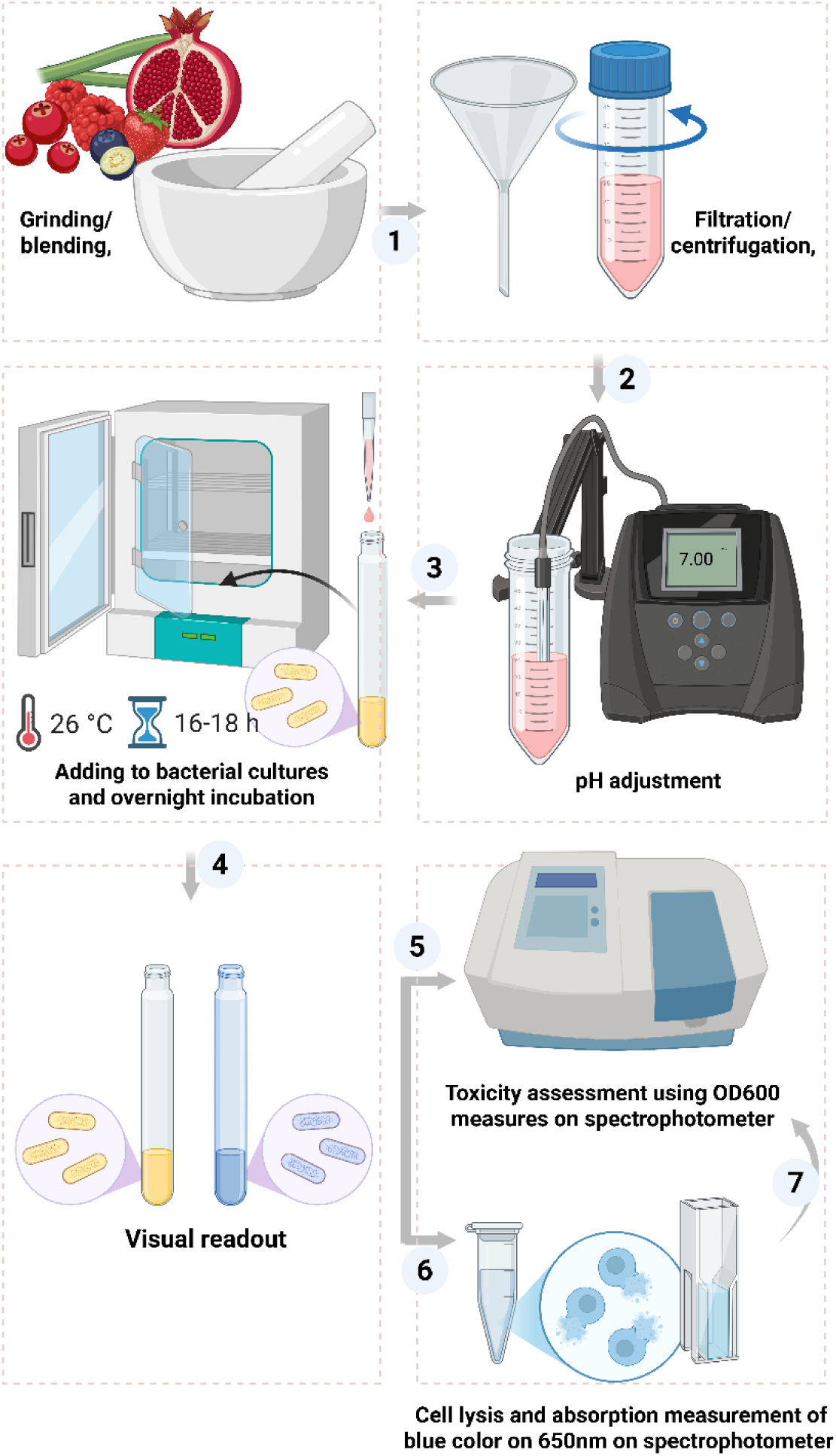
A schematic representation of experimental workflow summarizing the steps starting from 1) preparation of plant juices, up to 7) quantification of color development.

### 2.9. Statistical analysis

All samples were tested in triplicates. Given the small sample size (n = 3), no assumptions of normal distribution were made; therefore, a non-parametric Mann–Whitney U test was applied to compare independent groups. Statistical significance was accepted at p < 0.05. Calculations were performed in Microsoft Excel. Data were normalized to the “tested-juice-only” control (no X-gal) (see Table 1).

## 3. Results

### 3.1. Verification of the assembled genetic components of the system

Following correction of all detected sequence errors in the assembled, codon-optimized *Mpro* gene by iterative site-directed mutagenesis, agarose gel electrophoresis confirmed the expected amplicon size of approximately 921 bp (Figure 4). Sanger sequencing verified the integrity of the open reading frame, revealing no frameshift or nonsense mutations.

**Figure 4.**
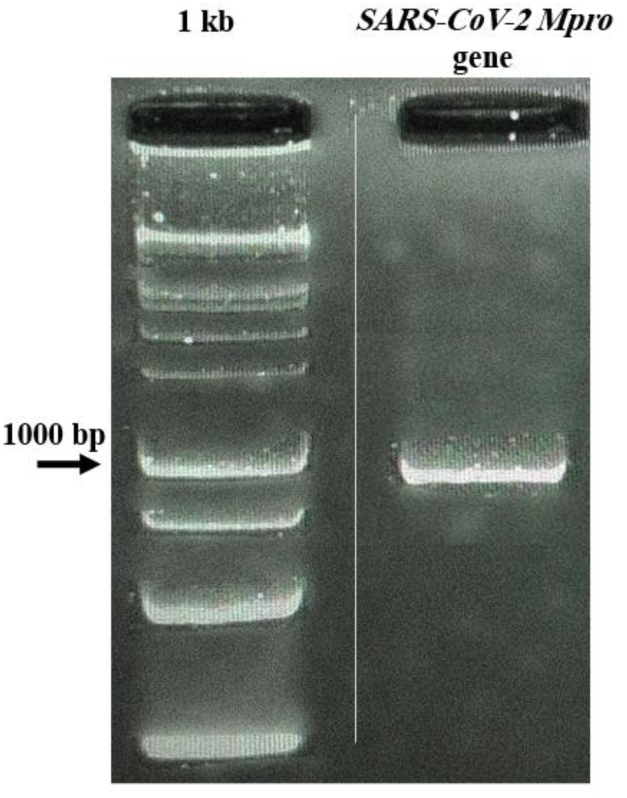
Gel electrophoresis for verification of assembled *Mpro* gene. A distinct band appears at approximately 921 bp, confirming full size of assembled gene.

Restriction digestion analysis confirmed the correctness of the final recombinant plasmid containing both the *Mpro* and modified *lacZα* genes. Sequencing further verified that insertion of the Mpro cleavage site preserved the *lacZα* reading frame. The complete construct comprising the lac promoter/operator-*Mpro*-T7 terminator cassette and the modified *lacZα* reporter was successfully assembled. A schematic plasmid map generated using Benchling software (Benchling Inc., USA) is presented in Figure 5. The verified pUC19-*Mpro* construct was used in all subsequent functional assays.

**Figure 5.**
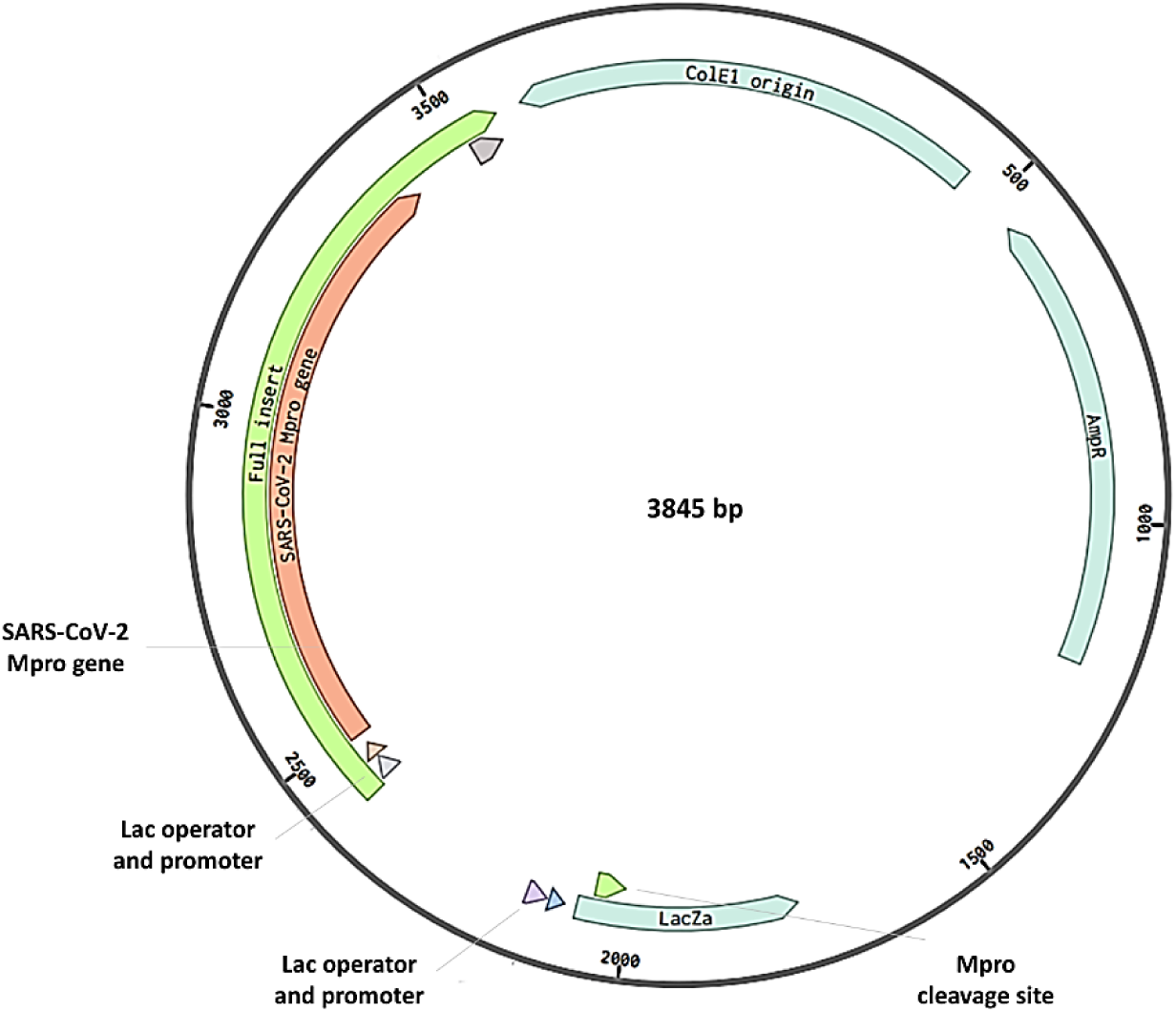
Map of constructed plasmid for the developed colorimetric reporter system, showing both genes; *Mpro* and reporter gene with inserted cleavage site of Mpro, under the same regulatory elements.

### 3.2. Expression validation of Mpro protein in E. coli DH5α

SDS–PAGE analysis comparing total cell lysates from *E. coli* DH5α carrying the recombinant pUC19–*Mpro* plasmid and control (cells with empty pUC19 vector), showed an additional distinct protein band in the recombinant sample, absent in the control. The apparent molecular weight of this band (∼33–34 kDa) corresponded to the expected size of SARS-CoV-2 Mpro (33.8 kDa), confirming successful expression under the applied induction conditions (Figure 6). This result verified that the loss of blue coloration observed in cells carrying both the *Mpro* and modified *lacZα* genes was indeed caused by the expression of functional Mpro.

**Figure 6.**
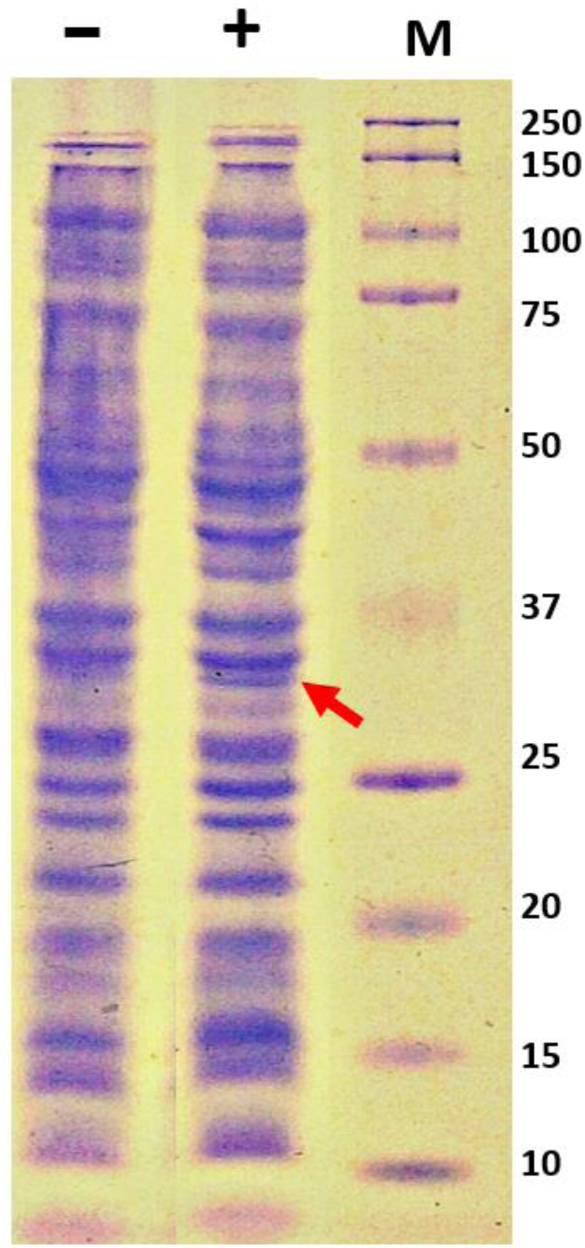
SDS–PAGE analysis of Mpro expression in recombinant *E. coli*. Lanes represent: (ـ) control strain lacking Mpro; (+) recombinant strain expressing Mpro; (M) Molecular weight marker; kDa (Bio-Rad Kaleidoscope™).

### 3.3. Functional validation of the system in living cells

The functionality of the constructed colorimetric reporter system was verified by comparing the phenotypes of *E. coli* DH5α cultures expressing both *Mpro* and the modified *lacZα* reporter with those expressing only the modified reporter. Following induction with IPTG in the presence of X-gal, cultures carrying only the reporter developed a vivid blue coloration, confirming intact β-galactosidase activity. In contrast, cultures co-expressing Mpro remained uncolored, indicating that the protease successfully cleaved its recognition site within *lacZα*, thereby disrupting β-galactosidase function (See 1 and 2 in **Figure 8**). These results confirmed that the system operated as designed and produced a distinct, visually detectable, and quantifiable phenotypic output suitable for inhibitor screening.

**Figure 7.**
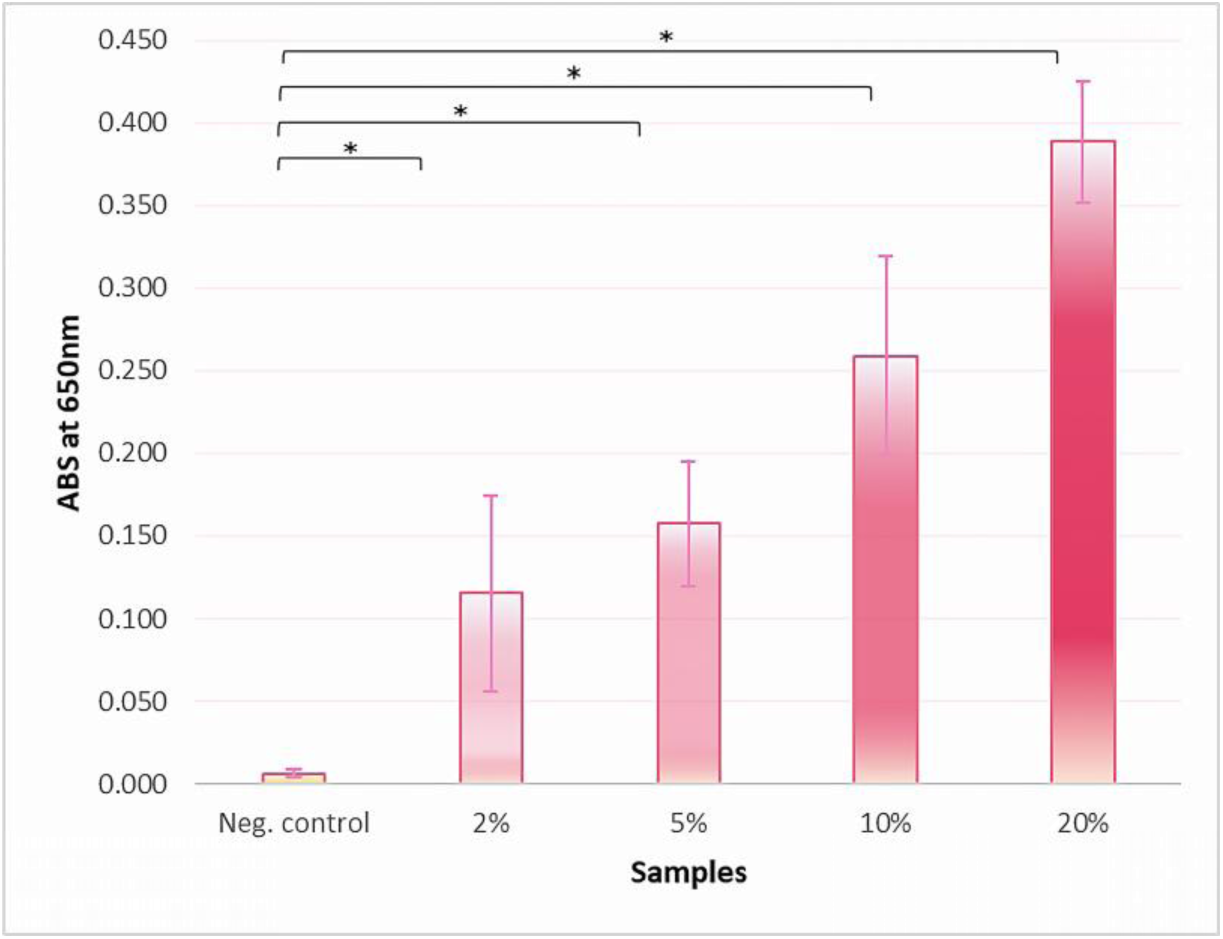
Concentration-dependent inhibition of Mpro by pomegranate juice. Bars represent mean absorbance at 650 nm ± SD. Asterisks indicate statistically significant differences in comparison with untreated control (p < 0.05).

**Figure 8.**
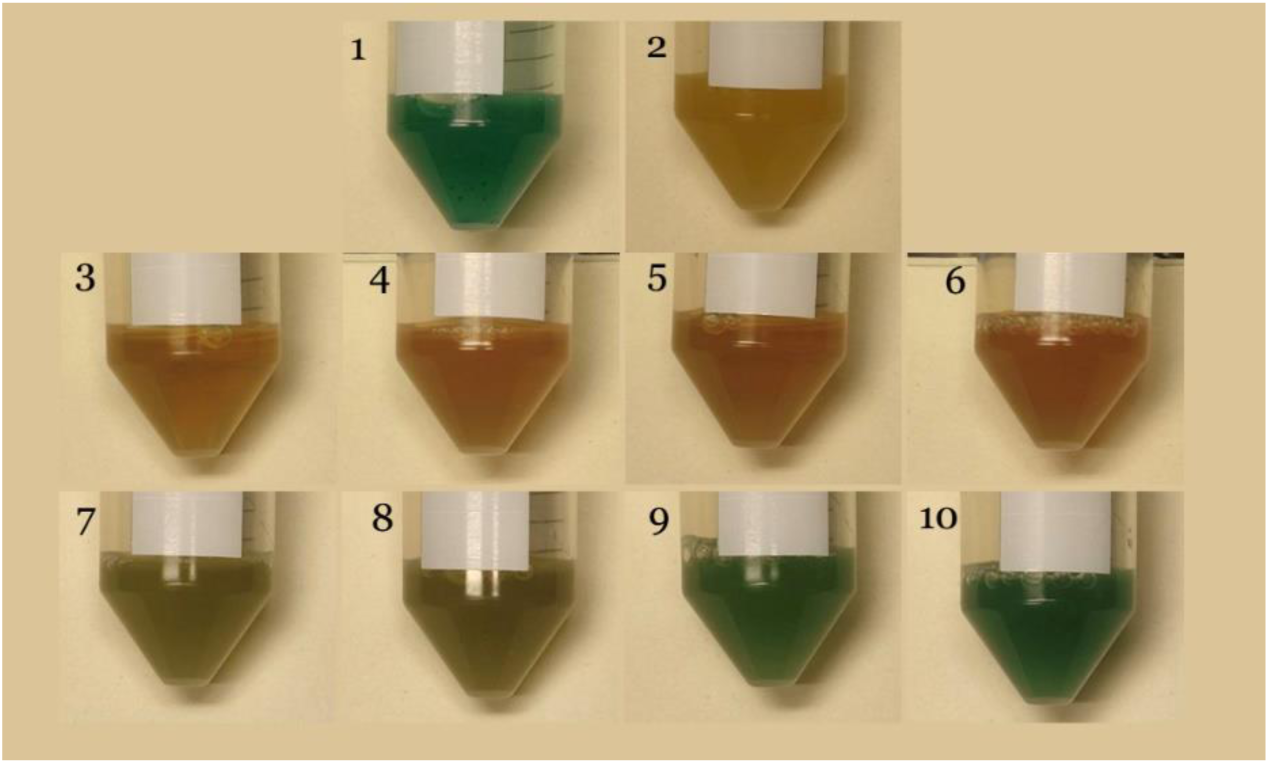
Visual readouts of controls and samples treated with pomegranate juice. 1) Reporter-only control, 2) untreated control, 3-6) Tested-juice-only control (no X-gal) in ascending concentrations (2%, 5%, 10%, and 20%, respectively), 7-10) samples treated with ascending concentrations (2%, 5%, 10%, and 20%, respectively) of pomegranate juice + X-gal.

### 3.4. Proof-of-concept validation of the screening system using pomegranate and guelder rose juices

For cultures incubated with pomegranate juice, absorbance at 650 nm, demonstrated a significant, concentration-dependent increase in blue coloration at all tested concentrations (p = 0.000) compared with their respective untreated controls, indicating inhibition of Mpro activity within living cells. The relative blue signal intensities (A650 values normalized to the control containing juice without X-gal) are presented in Figure 7, and visual readouts are shown in Figure 8.

Assessment of optical density (OD_600_) before and after incubation with pomegranate juice showed a significant concentration-dependent reduction in growth at concentrations (5%, 10%, and 20%), compared to untreated control (Figure 9). Despite this growth inhibition, blue coloration remained clearly detectable, indicating that the residual viable cell population retained sufficient β-galactosidase activity for the system to generate a positive colorimetric readout.

**Figure 9.**
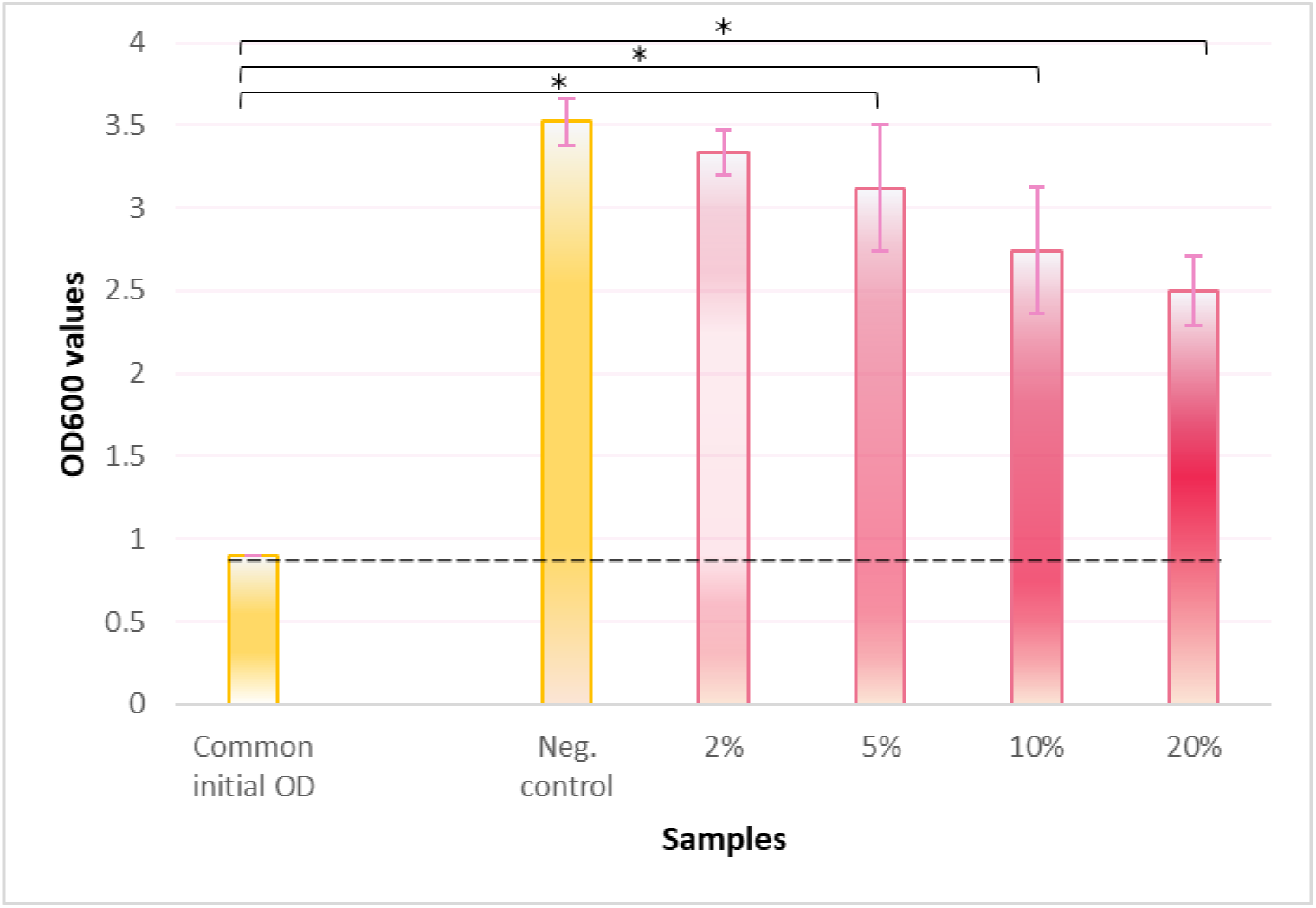
OD_600_ measurements before and after pomegranate juice treatment. Initial OD_600_ before induction was standardized. Asterisks indicate statistically significant differences in comparison with untreated control (p < 0.05).

In contrast, cultures incubated with guelder rose juice also exhibited strong and consistent growth inhibition across tested concentrations (significant reduction in OD_600_ values), however, no increase in blue coloration was observed at any concentration. Because the cell population was substantially reduced and no measurable color development occurred, the juice was excluded from quantitative analysis.

These results confirm the ability of the developed *E. coli*-based system to detect intracellular Mpro inhibition using plant preparations even in conditions of partial growth inhibition/mild toxicity.

### 3.5. Internal controls validation and assay reliability

Performance of the set of internal controls included to ensure the assay’s reliability is summarized in Table 2.

**Table 2.** A summary of observed outcomes for all internal controls.

| Control | Observed outcome |
| --- | --- |
| LB-only control | No signal or growth, indicating a clean background. |
| Reporter-only control | Strong blue coloration, confirming β-galactosidase activity (positive baseline). |
| Vehicle control | No effect on color or growth, confirming solvent neutrality. |
| Untreated control | No coloration observed with X-gal, confirming Mpro activity. |
| Tested-juice-only control (no X-gal) | Only native plant pigment observed; no blue coloration (negative baseline). |
| Reporter-only + Plant juice control | Blue coloration mixed with natural pigment, serving as color reference for positive inhibition effects. |

## 4. Discussion

This study demonstrates the successful construction and validation of an *E. coli*–based colorimetric reporter system for detecting inhibition of the SARS-CoV-2 Mpro directly in living cells by plant crude preparations (juices) and other potential inhibitors. The developed system links Mpro functionality to a quantifiable visual phenotypic output, enabling assessment of protease inhibition by tested juices in a safe, simple, and cost-effective way. Such a tool offers a practical alternative to existing computational, biochemical, and eukaryotic models, particularly valuable during early-stage antiviral screening under limited laboratory conditions.

The correctness and integrity of the assembled genetic components were first validated by sequencing and restriction analyses, confirming that the codon-optimized *Mpro* gene and the modified *lacZα* reporter were correctly constructed and maintained in frame. This step was essential to ensure the functional linkage between both expressed proteins, and, thereby, between Mpro enzymatic activity and the resulting phenotypic output. Sequence errors introduced during synthesis and amplification were corrected by site-directed mutagenesis, while codon optimization enhanced translational efficiency without altering the encoded amino acid sequence. Such strategies have been reported to enhance heterologous viral protein production in bacterial systems, supporting the reliability of this design for intracellular bacterial expression of SARS-CoV-2 Mpro [23]. Expression analysis confirmed the production of an intact Mpro protein with an apparent molecular mass consistent with its theoretical size of 33.8 kDa. Detection of this band in DH5α lysates indicated that even a cloning host strain, which is not optimized for protein overexpression, can produce sufficient protease levels for measurable activity, confirming the efficiency of the construct. The observed loss of blue coloration upon co-expression of both genes, verified that Mpro reserved its catalytic activity in the bacterial cytoplasm, and specifically cleaved its specific site within the modified *lacZα* gene product. This demonstrated that Mpro was not only expressed but also functionally active, providing further evidence of the construct’s efficiency.

The obtained binary color change: blue coloration in the absence of protease activity and its loss when Mpro was expressed, provided both qualitative and quantitative output suitable for screening. In contrast with *in silico*, biochemical, and mammalian cell-based assays, this system represents an efficient middle ground. *In silico* screening, while cost-effective, can only predict or filter candidates without accounting for cellular uptake, solubility, or toxicity. Biochemical assays typically require purified protein and specialized instrumentation, while mammalian systems, despite physiological relevance, are costly, less safe, and demand advanced biosafety infrastructure. The *E. coli*–based system simultaneously reflects compound stability and intracellular bioavailability while remaining experimentally simple and accessible.

Proof-of-concept validation using pomegranate (*Punica granatum*) juice confirmed the system’s responsiveness to natural inhibitors. A significant increase in blue coloration at higher concentrations indicated successful inhibition of Mpro activity within bacterial cells. Pomegranate was selected due to its reported strong antiviral, antioxidant, and polyphenol-rich profile, as well as several studies demonstrating inhibition of SARS-CoV-2 Mpro by its ellagitannins and punicalagins [24, 25]. Similarly, guelder rose (*Viburnum opulus*) was included because of its high polyphenolic content and its traditional medicinal use, particularly in Eastern Europe and Russia, where it is widely applied against respiratory conditions [26]. Juices, rather than purified extracts, were used because they offered the simplest test material without the need for extraction, further demonstrating the system’s simplicity. Importantly, juices also allowed assessment of potential synergistic effects among multiple phytochemicals, rather than isolated compounds. From a nutritional and practical perspective, evaluating dietary juices was relevant, as the focus in urgent pandemic situations should prioritize safe, accessible, and naturally derived antiviral sources [19]. The tested concentration range was selected to represent moderate exposure levels; sufficient to deliver active compounds while avoiding excessive sugar or polyphenol content that could affect bacterial physiology or optical readings. Lower concentrations (<2%) were excluded because intracellular exposure to active compounds would likely fall below the detection threshold, while higher concentrations (>20%) could increase medium turbidity, viscosity, or nonspecific metabolic inhibition. Strong pH adjustment was avoided to examine potential effects of naturally occurring acids (while still relatively preserving bacterial tolerance to acidity), such as ascorbic, caffeic, chlorogenic, ellagic, ferulic, gallic, protocatechuic, and ursolic acids [27, 28], many of which have been reported to inhibit SARS-CoV-2 Mpro [24, 29]. The concentration-dependent restoration of blue coloration observed for pomegranate supported its suitability as a proof-of-concept inhibitor source. Notably, despite the concentration-dependent growth suppression detected in OD_600_ measurements, the system still produced a clear colorimetric response at higher concentrations, demonstrating that it remains functional even under partial growth inhibition. In contrast, guelder rose caused strong growth suppression across tested concentrations but did not generate detectable color restoration, which is consistent with a lower or absent content of active Mpro-targeting phytochemicals at the tested concentrations. Importantly, this differential outcome highlights the system’s ability to distinguish active from inactive plant crude preparations, as only juices with functional Mpro-inhibitory activity result in restoration of the blue phenotype, while unrelated phytochemical profiles do not generate false-positive signals. Additionally, this shows that reduced cell viability does not result in false-negative inhibition outcomes.

The comprehensive set of internal controls verified assay specificity and robustness by ensuring the validity and interpretability of the visual outcome. These included treated reporter-only controls to assess assay responsiveness, reporter-only and untreated controls to confirm baseline activity and reporter integrity, and juice-only, vehicle, and contamination controls to rule out non-specific color or growth effects. Together, these controls establish clear interpretive baselines for distinguishing true inhibitory activity from unrelated visual or metabolic changes.

During optimization, several technical challenges were encountered and systematically resolved. Initially, a dual-plasmid configuration was tested, in which *E. coli* cells were co-transformed with one plasmid encoding SARS-CoV-2 Mpro under the lac promoter/operator and another containing the *lacZ* reporter gene interrupted by the engineered Mpro cleavage site. However, unbalanced gene expression, unequal plasmid copy-numbers, and potential incompatibility of replication origins introduced substantial variability in color development, complicating interpretation. Transition to a single-plasmid construct unified gene regulation and eliminated variability. Additionally, early assays were performed on solid LB-agar supplemented with IPTG, X-gal, and ampicillin, however, yielded uneven color distribution and inconsistent quantification, which were resolved by adopting liquid cultures that ensured uniform distribution and allowed quantitative spectrophotometric assessment. Induction at higher cell densities (OD_600_ = 0.6–0.9) enhanced assay sensitivity toward weaker inhibitory effects, whereas lower induction ODs produced less distinct color differentiation. IPTG concentrations of 0.1–0.5 mM and X-gal concentrations below 0.2 mg/mL were evaluated; the selected combination of 0.2 mM IPTG and 0.2 mg/mL X-gal offered optimal balance between induction efficiency and visual contrast. Temperature optimization revealed that incubation at 26 °C provided the most consistent color development with minimal growth inhibition. Additionally, several buffer formulations were tested for efficient cell lysis without altering the chromogenic pigment. The final buffer composition achieved reproducible solubilization and color stability. These troubleshooting steps collectively enhanced assay reproducibility and demonstrated the system’s adaptability to complex sample types, ensuring optimal assay performance.

As mentioned above, the developed system presents several key advantages. It assesses inhibition in living cells, thereby accounting for protein expression, folding, and stability. Its gain-of-signal design minimizes false negatives, since color restoration reflects inhibition rather than signal loss, which in other assays could result from toxicity or metabolic disruption. The assay is rapid, simple, and compatible with standard laboratory equipment, avoiding the need for advanced instrumentation or costly eukaryotic models. It is also biosafe and noninfectious, as it does not involve live virus handling, and allows straightforward interpretation and quantification. The system also enables built-in monitoring of cytotoxicity through bacterial growth observations. Importantly, it has been proven capable of testing complex plant preparations, not only purified compounds, supporting its use in evaluating synergistic and cumulative effects of dietary plants as potential preventive or supportive measures against viral infections.

Despite these strengths, several limitations should be acknowledged. The *E. coli* outer membrane restricts permeability to large molecules, limiting the system’s ability to assess macromolecular or poorly permeable inhibitors such as nirmatrelvir, which could otherwise serve as a positive control for Mpro inhibition in the present study. The bacterial cytoplasmic environment also differs from that of mammalian cells, so compounds requiring metabolic activation or specific transporters may not behave identically. Furthermore, highly colored or viscous extracts could interfere with optical readings at extreme concentrations, although internal normalization largely compensates for this. Therefore, the system is not intended to replace high-resolution biochemical or antiviral efficacy assays but rather to serve as a preliminary, prioritization-stage screening tool. Candidate hits from this system can subsequently be validated in mammalian or biochemical models. A concise comparison of the developed system with existing approaches is provided in Table 3, summarizing their main features, advantages, and limitations.

| Approach | Key features | Advantages | limitations |
| --- | --- | --- | --- |
| In silico<br>(computational docking, molecular dynamics) | Virtual prediction of binding affinities based on Mpro structure | Fast, inexpensive, suitable for large-scale screening | Lacks biological context; no data on cell permeability, solubility, or toxicity |
| Biochemical<br>(cell-free<br>enzyme assays,<br>fluorescence/luminescence<br>readouts) | Recombinant purified<br>Mpro tested with<br>substrates in vitro | Quantitative, high<br>specificity, allows<br>kinetic analysis | Requires protein<br>purification and<br>specialized<br>equipment,<br>ignores cellular<br>uptake |
| E. coli-based<br>colorimetric<br>reporter system<br>(present study) | Mpro activity linked to β-<br>galactosidase reporter<br>within living bacterial<br>cells | Detects inhibition<br>in a cellular<br>environment;<br>accounts for<br>compound<br>stability, and<br>bioavailability,<br>simple visual and<br>spectrophotometric<br>quantification,<br>cost-effective,<br>biosafe, adaptable<br>for large-scale<br>screening and for<br>other<br>proteases/mutants | Limited by<br>bacterial<br>permeability and<br>potential<br>interference from<br>highly colored<br>extracts, not fully<br>representative of<br>mammalian<br>metabolism |
| Mammalian<br>cell-based<br>models | Mpro expression or viral<br>replication assays in<br>human cell lines | Physiologically<br>relevant,<br>integrates host<br>metabolism | Expensive, time-<br>consuming,<br>requires high-<br>biosafety facilities<br>and expertise,<br>potential safety<br>risks |

In a broader perspective, beyond SARS-CoV-2 Mpro, the modular design can be readily adapted to other viral or bacterial proteases/mutant variants by substituting their specific cleavage motifs. It can also be optimized for high-throughput applications, enabling faster primary screening. The low biosafety requirements, minimal instrumentation needs, and visual readout make it particularly suitable for preliminary research or educational settings where advanced cell culture facilities are unavailable. Future work may also include the use of membrane-permeabilizing controls to expand the range of testable compounds.

## 5. Conclusions

Overall, the results confirm that the developed system successfully couples protease activity to a visible phenotypic change, providing a reliable, and accessible approach for identifying potential protease inhibitors from natural sources. It represents a practical and biosafe tool for preliminary evaluation of natural Mpro inhibitors and can be adapted for broader antiviral screening purposes.

## Author Contributions

Conceptualization, T.V.M.; methodology, S.S.I., A.A.Z., H.J.F., R.R.Z., and T.V.M.; validation, S.S.I., H.J.F., and T.V.M.; formal analysis S.S.I., R.R.Z., and T.V.M.; investigation, S.S.I., H.J.F., and T.V.M.; resources, A.A.Z. and T.V.M, writing—original draft preparation, S.S.I., H.J.F., and T.V.M.; writing—review and editing, S.S.I. and T.V.M.; visualization, T.V.M.; supervision, T.V.M.; project administration, T.V.M. All authors have read and agreed to the published version of the manuscript.

## Acknowledgments

The work was performed using the equipment of the Resource Center of Saint Petersburg State University “Development of Molecular and Cellular Technologies”.

## Abbreviations

The following abbreviations are used in this manuscript:

SARS-CoV-2 Mpro, 3CLpro, Nsp5: The main protease of novel coronavirus, or 3C-like protease, or non-structural protein 5
*E. Coli*: *Escherichia coli*
IPTG: isopropyl β-D-1-thiogalactopyranoside
v/v: Volume to volume ratio
OD_600_: Optical density at 600 nm
A650: Absorption at 650 nm

